# Interactions of cytosolic termini of the Jen1 monocarboxylate transporter are critical for trafficking, transport activity and endocytosis

**DOI:** 10.1101/2021.09.27.461913

**Authors:** Cláudia Barata-Antunes, Gabriel Talaia, George Broutzakis, David Ribas, Pieter De Beule, Margarida Casal, Christopher J. Stefan, George Diallinas, Sandra Paiva

## Abstract

Plasma membrane (PM) transporters of the major facilitator superfamily (MFS) are essential for cell metabolism and growth, as well as for survival in response to stress or cytotoxic drugs, in both prokaryotes and eukaryotes. In the yeast *Saccharomyces cerevisiae*, Jen1 is a monocarboxylate/H^+^ symporter that has been used to dissect the molecular details underlying control of cellular expression, transport mechanism and turnover of MFS transporters. Here, we present evidence supporting previously non-described roles of the cytosolic N- and C- termini in Jen1 biogenesis, PM stability and activity, through functional analyses of rationally designed truncations and chimeric constructs with UapA, a *S. cerevisiae* endocytosis-insensitive purine transporter from *Aspergillus nidulans*. Our results reveal a cryptic role of the N-terminal region and thus show that both cytosolic N- and C-termini are critical for Jen1 trafficking to the PM, transport activity and endocytosis. In particular, we provide evidence that the N- and the C-cytosolic termini of Jen1 undergo transport-dependent dynamic intra-molecular interactions, which critically affect the mechanism of transport and turnover of Jen1. Our results support an emerging concept where the cytosolic tails of PM transporters control transporter expression and function, through flexible intra-molecular interactions with each other and the transmembrane core of the protein. This idea may be extended to other MFS members providing a deeper understanding of conserved, but also evolving, mechanisms underlying MFS transporter structure-function relationships.

## Introduction

Eukaryotic plasma membrane (PM) transporters play essential roles in cell nutrition, signalling, and responses to stress conditions and drugs. Consequently, transporter malfunction has an impact in many aspects of human cell biology and leads to several pathologies, including neurological and cardiovascular disorders, as well as diabetes and cancer (1–5). Given their importance in sensing the environment and maintaining cell homeostasis, transporter function depends on complex and fine regulatory mechanisms. Endocytic internalization is a major regulatory mechanism of PM transporters, mostly studied in the model fungi *Saccharomyces cerevisiae* and *Aspergillus nidulans*, in response to physiological or stress signals, followed by either their vacuolar degradation or recycling back to the PM via the TGN/endosomal system (for recent reviews see (6–8)). Endocytic internalization of fungal transporters requires ubiquitylation at their C- or N-terminal cytosolic regions by HECT-type E3 ubiquitin ligases (e.g., Rsp5 in *S. cerevisiae* or HulA in *A. nidulans*), which are recruited by adaptor proteins named α-arrestins (9–16). In *S. cerevisiae*, fourteen α-arrestins have been identified, named Arts (Art1-10), Buls (Bul1-3) and Spo23, which all possess PY motif(s) that may interact with WW domains of Rsp5 Ub ligase, mediating membrane protein turnover (9, 10, 12, 17–19). *A. nidulans* possesses 10 α-arrestins, including ArtA and PalF, which control transporter down-regulation and sensing, respectively (8, 14). In mammals, six α-arrestins have been identified, named ARRDC proteins (20, 21), but much less is known regarding their role, specifically on transporter cellular expression. Notably, however, ARRDC6/TXNIP has been shown to function as an endocytic adaptor for the GLUT1 and GLUT4 transporters (22, 23).

To exert their function, α-arrestins need to recognize the cytoplasmic exposed segments of transporters, basically their N- or C-termini. Pioneering studies in *S. cerevisiae* have shown that the N-terminus parts of amino acid transporters Can1 and Lyp1 are specifically recognized by α-arrestins Art1 and Art2, respectively, under stress conditions (9). The N-terminus of the general amino acid transporter Gap1 also contains a potential Bul1/2 α-arrestin interacting motif and two Ub target sites (K9 and K16) that are required for nitrogen-elicited endocytosis of Gap1 (24, 25). Additionally, under stress conditions, Bul1/2, in combination with Art3/Art6, promotes Gap1 ubiquitylation and down-regulation *via* Gap1 C-terminus (26). In these cases, it is thought that conformational changes during substrate transport makes the N-terminus more accessible to α-arrestins (25, 27). The methionine-specific transporter Mup1 also possess in its N-terminus a motif, proximal to the ubiquitylation sites (K27 and K28), which is proposed to act as a putative Art1 α-arrestin target site required for substrate-elicited ubiquitylation and endocytosis (28). A putative Art1 interacting motif has also been found for Can1 (27, 28). In fact, Can1 N-terminus possesses specific lysines and two putative α-arrestin interacting motifs (Art1 and Bul1/2), which are required for substrate-elicited ubiquitylation and endocytosis of the permease (27). The Fur4 uracil transporter N-terminus possesses Ub acceptor sites (K31 and K41) involved in ubiquitylation and endocytosis (29–31). Noticeably, the Fur4 N-terminus is likely to undergo dynamic conformational changes, in response to excess of substrate or stress, enhancing its endocytic down-regulation (31).

In the filamentous fungus *A. nidulans,* a C-terminus region of the uric acid transporter UapA is essential for ArtA-mediated ubiquitylation, endocytosis and vacuolar degradation in response to ammonium or excess of substrate (11, 14). *A. nidulans* Fur4 homologues have also been shown to possess elements in their N- and C-terminus that are critical for endocytosis and surprisingly substrate specificity. In this case, the authors provided evidence that the N- and the C-terminus interact physically and promote proper transporter function and turnover (32, 33).

Whether long-range regulatory effects of cytosolic N- and C-termini extend to transporters other than Fur-like proteins and a handful of other members of the amino acid–polyamine–organocation (APC) superfamily (34), remains to be formally shown.

Here, we address this issue by using Jen1, a well-studied yeast transporter that represents the ubiquitous and largest transporter family, namely the major facilitator superfamily (MFS). In particular, we genetically and functionally dissect the role of both cytosolic N- and C-termini of Jen1 and provide compelling evidence for a cryptic role of the N-terminus of Jen1, which together with sequence elements in the C-terminal region, control the biogenesis, activity and turnover of Jen1. Most importantly, using quantitative bi-fluorescence complementation (BiFC) assays, we present evidence that the two Jen1 termini interact dynamically in a transport-activity dependent manner, which ultimately regulates Jen1 cell-surface expression and activity. Our findings support the idea that cytosolic tails in eukaryotic transporters have acquired important multi-functional roles and thus reveal novel regulatory mechanisms of MFS family members.

## Results

### Rationale for constructing specific Jen1 truncations and chimeric transporters

Jen1 is a specific monocarboxylate/H^+^ symporter (lactate and pyruvate being its major substrates) that has been used extensively as a model cargo to dissect mechanisms of regulated transporter internalization. Jen1 ubiquitylation, endocytosis and vacuolar degradation are regulated by two α-arrestins (Rod1 and Bul1), in response to distinct stimuli (16, 35). Rod1-mediated endocytosis of Jen1 requires the presence of a preferred carbon source, such as glucose, in a substrate transport-independent manner (16, 35). In addition, conformational changes associated to substrate transport are likely to trigger Bul1-mediated endocytosis of Jen1, in response to alkali stress (16). Recently, a C-terminal region of Jen1 was reported to be involved in Rod1-mediated endocytosis of the transporter, triggered by glucose (36). Several specific lysines in Jen1 protein were reported to be required for its ubiquitylation and endocytosis (15, 35–37).

To address the role of the cytosolic termini of Jen1 in its regulation, we employed specific N- or C-terminus truncations of Jen1, as well as chimeric transporters based on the UapA transporter from *A. nidulans*, carrying the cytosolic termini of Jen1. UapA is an extensively studied uric acid-xanthine/H^+^ symporter (for a review see (38)), which is regulated by ammonium or substrate-elicited endocytosis in *A. nidulans*. However, upon functional expression in *S. cerevisiae*, it does not respond to endocytosis and, instead, remains stable at the PM (39). Thus, UapA provides an appropriate molecular marker for investigating, via domain swap experiments, the potential, context-independent, functional role of *cis*-acting elements present in Jen1 N- or C-terminal regions.

Prior to these constructions, it was essential to define the limits of the N- and C-terminus of Jen1 based on available structural information. The selection of the number of residues corresponding to the cytosolic N- and C-terminus portions of Jen1, which lacks an experimentally defined structure, was based on standard topology predictions and homology threading modelling, using various bioinformatic tools (detailed in **Table S1**). These predictions were used to construct three Jen1 truncated versions by deleting the longest predicted N-terminal region (133 residues) and the two versions of the putative C-terminus (62 or 33 residues) of Jen1. The resulting truncated versions were named Jen1ΔNT133, Jen1ΔCT62 or Jen1ΔCT33 (**Figure 1A**). Based on the recent work of Fujita and co-workers (2018), we also generated a shorter N-terminal truncation (Jen1ΔNT94) and the doubly truncation Jen1ΔNT94ΔCT33. Chimeras of UapA/Jen1 were constructed as illustrated in **Figure 1B**. Briefly, the intact UapA sequence was fused with amino acid segments 1-94 or/and 584-616 of Jen1 termini, resulting in the chimeric transporters named UapA/Jen1NT94, UapA/Jen1CT33 and UapA/Jen1NT94-CT33.

**Figure 1.**
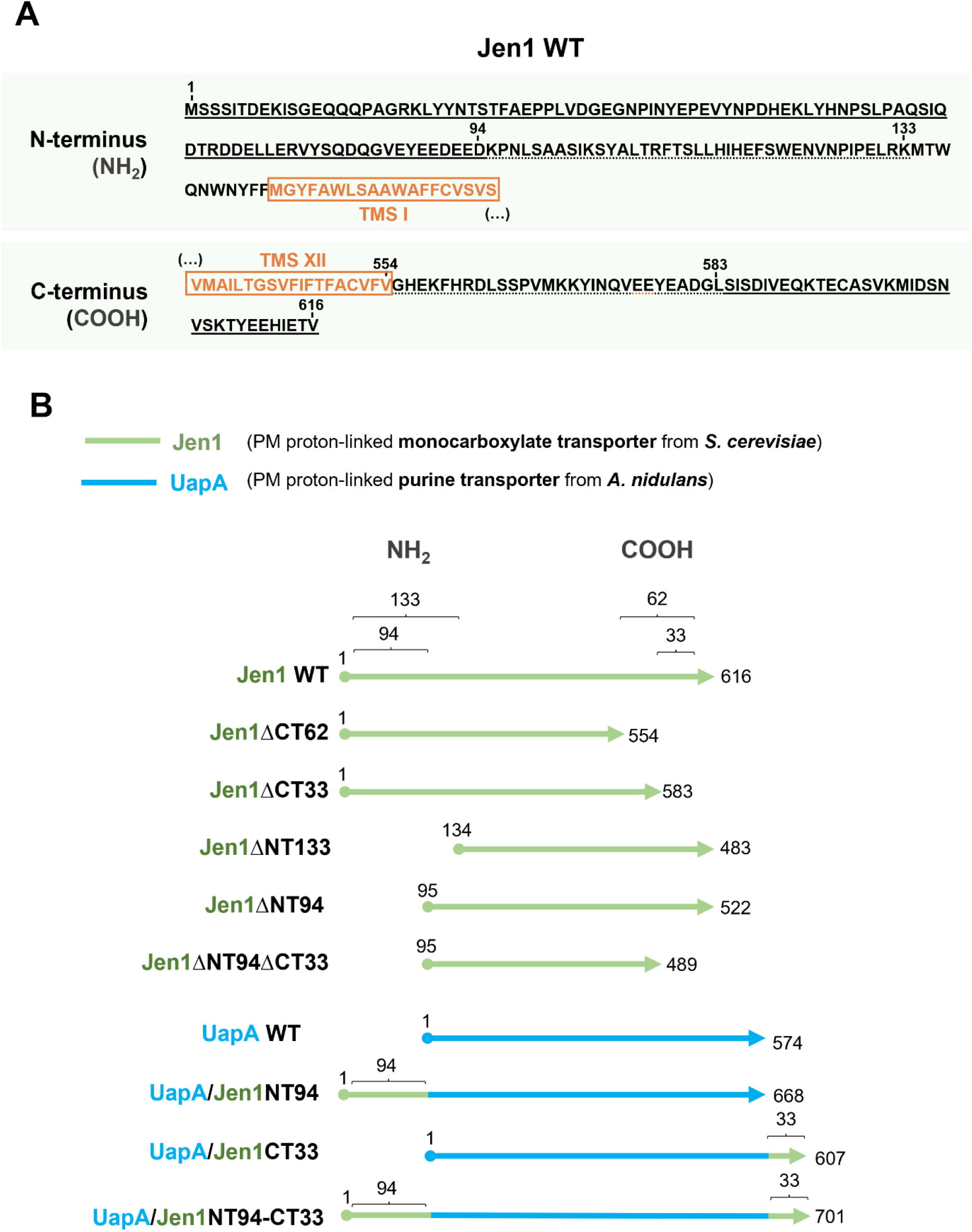
Designing of truncations and chimeric versions of Jen1. **(A) Primary amino acid sequences of the N- and C-terminal regions of Jen1 transporter.** N- and C-terminal predicted regions of Jen1 transporter are shown in black letters. The number of residues corresponding to these regions is indicated. Residues framed in orange correspond to predicted first or last transmembrane segments (TMSI and TMSXII, respectively), as defined by secondary prediction programs (see in Experimental Procedures). **(B) Graphical representation of truncated Jen1 versions and Jen1-UapA chimeric transporters.** Jen1 and UapA sequences are shown in green and blue, respectively. Jen1 mutant versions were cloned either under the control of the strong GPD (glyceraldehyde-3-phosphate dehydrogenase) promoter, which allows the constitutive expression of *JEN1* (40), or under the control of the *GAL* promoter, enabling the expression of *JEN1*, under galactose (2 %, w/v) inducible conditions.

### Specific N- and C-terminally-truncations result in Jen1 versions with modified PM localization, stability and transport kinetics

We analysed the growth pattern and transport activities of strains expressing Jen1 truncations, compared to those observed with a strain expressing wild-type Jen1 (**Figure 2A** and **2B**). In all cases, Jen1 versions were functionally tagged, C-terminally, with GFP. Phenotypic assays demonstrated that all shorter Jen1 termini truncations (Jen1ΔNT94, Jen1ΔCT33 and Jen1ΔNT94ΔCT33) were able to confer lactate growth (as sole carbon source) similar to the wild-type Jen1. In contrast, the longer cytosolic Jen1 truncations (Jen1ΔNT133 and Jen1ΔCT62) scored as *null* Jen1 mutants in growth tests (**Figure 2A**). Jen1-mediated lactate transport activity measurements, performed under Jen1-derepressed conditions (see Materials and methods), showed that Jen1ΔNT133 and Jen1ΔCT62 truncations displayed residual or no lactate transport activity, in line with growth tests. Also, in accordance with growth tests, Jen1ΔNT94, Jen1ΔCT33 and Jen1ΔNT94ΔCT33 truncations were able to import lactate with apparent rates similar to those measured in the wild-type Jen1 (**Figure 2B**). The recorded transport capacities in the mutants were in agreement with measurements of alkalinization of the external medium via Jen1-dependent lactate uptake (16), as among the truncations, only Jen1ΔNT94, Jen1ΔCT33 and Jen1ΔNT94ΔCT33 led to an increase in the pH of the medium (**Figure S1A**).

**Figure 2.**
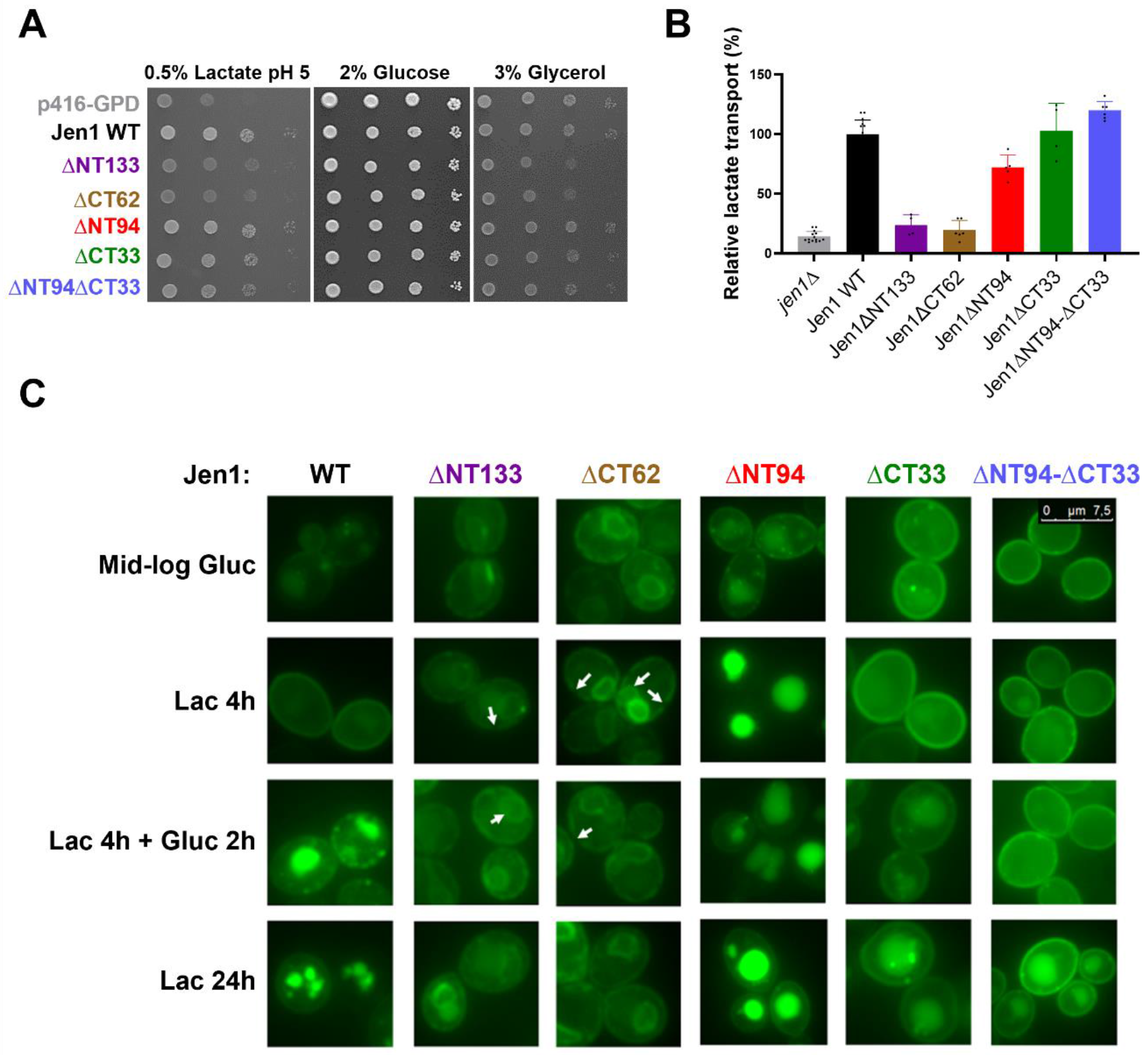
Removing specific segments of Jen1 cytoplasmic N- and C-termini modifies protein PM localization and transport kinetics. *S. cerevisiae jen1*Δ cells expressing the empty p416-GPD plasmid (*jen1*Δ), the *JEN1* gene (Jen1 WT), or five *JEN1* mutant versions (Jen1ΔCT33, Jen1ΔCT62, Jen1ΔNT133, Jen1ΔNT94 and Jen1ΔNT94ΔCT33), tagged with GFP, were characterized by growth assays (**A**), transport uptakes (**B**) and epifluorescence microscopy (**C**). (**A**) Serial 1:10 dilutions of yeast cells were spotted onto YNB containing plates supplemented with three distinct carbon sources: glucose (2 %, w/v), glycerol (3 %, v/v) or lactate (0.5 %, v/v, pH 5.0). Cells were grown for 7 days at 18 ⁰C. (**B**) Percentage of ^14^C-lactic acid uptake, at pH 5.0, in YNB lactic acid-derepressed cells. The rate of wild-type Jen1 is taken as 100 %. Individual data points are shown. Error bars correspond to standard deviation values. (**C**) Epifluorescence microscopy analysis of Jen1-GFP or its derivatives. Samples were collected after growth on glucose (Mid-log Gluc), after derepression in lactate medium for 4 hours (Lac 4 h), after 2 hours of a pulse of glucose (2 %, w/v) to Lac 4 h induced cells (Lac 4 h + Gluc 2 h) or after prolonged growth on lactate (Lac 24 h).

The subcellular localization of Jen1 truncations (**Figure 2C**) was followed under conditions that promote Jen1 localization to the PM (Lac 4 h), or conditions that lead to endocytic turnover, such as addition of glucose (Lac 4 h + Gluc 2 h) or prolonged growth on lactate (Lac 24 h) (16). As expected, stable localization of wild-type (WT) Jen1 to the PM was observed upon lactate induction (Lac 4 h), while endocytosis and sorting to vacuoles for degradation was observed upon 2 h of growth in the presence of glucose, and more dramatically after 24 h growth on lactate. The larger truncations Jen1ΔNT133 and Jen1ΔCT62 showed significant ER retention of the protein, revealed by fluorescent labelling of perinuclear ER rings, but also discontinuity of the fluorescence signal at the cell periphery, typical of cortical ER (cER) in yeast. Co-staining with CMAC excluded that the observed intracellular rings correspond to vacuoles (**Figure S1B**). In the case of Jen1ΔCT62, localization to the ER was formally confirmed by expression in a *S. cerevisiae* strain lacking all six ER-PM tethering proteins. In this strain (called Δtether), the cER has no contact with the PM so that ER resident proteins can be unambiguously distinguished from those localized to the PM (41). Using this strain, we showed that the cER, marked with an ER-resident red-fluorescing marker (DsRED-HDEL), co-localized fully with Jen1ΔCT62-GFP, but not with the functional truncation Jen1ΔCT33-GFP (**Figure S2A**).

The smaller functional Jen1 truncations gave a rather surprising result in respect to PM localization. Jen1ΔNT94, although being functional, as demonstrated by growth tests and transport uptakes (**Figure 2A** and **2B**), proved to be unstable in terms of subcellular localization, undergoing very rapid internalization and vacuolar degradation (**Figure 2C**, **Figure S1B**). This suggests that Jen1ΔNT94, after basal constitutive expression, is very sensitive to endocytosis in response to the presence of its substrate (lactate) or glucose. In sharp contrast, Jen1ΔCT33 homogeneously and stably localized to the PM with no indication of ER retention or vacuolar degradation after 4 h lactate induction (**Figure 2C**, **Figure S2A**). Notably, Jen1ΔCT33 led to a stronger fluorescence signal associated with PM compared to the wild-type, suggesting that this truncation stabilizes the transporter. This justified the moderate increase in the relative apparent transport activity obtained by direct uptake measurements (**Figure 2B**). Jen1ΔCT33 also showed reduced endocytosis upon glucose addition or after prolonged growth on lactate, the latter being more evident when compared to the wild-type control (**Figure 2C**, **Figure S1B**). Reduced endocytosis might well be due to the increased stability of this Jen1 version, observed in the absence of signals triggering endocytosis (i.e., 4 h Lac). Most surprisingly, the doubly truncated Jen1ΔNT94ΔCT33 version was also stably localized to the PM, similar to Jen1ΔCT33, and, in addition, it was more resistant to both signals triggering endocytosis, when compared to both WT Jen1 and Jen1ΔC33 (**Figure 2C**, **Figure S1B**). This suggested that truncating the 33 last residues of Jen1 is not just epistatic to the instability conferred by deleting the N-terminal 94 residues, but also pointed to the idea that the two termini of Jen1 interact functionally.

To better understand the combinatorial effect of deletions of terminal segments on Jen1 function, we investigated, via direct measurements of lactic acid transport, whether functional truncations (Jen1ΔNT94, Jen1ΔCT33 and Jen1ΔNT94ΔCT33) affect the transport kinetics of lactate (**Figure 3**). All Jen1 truncations tested, including Jen1ΔNT94, which proved to be an unstable version of Jen1, displayed higher substrate affinities (lower *K_m_*) compared to wild-type Jen1. Notably, deleting the C-terminal region in Jen1ΔCT33 resulted in a 10-fold increase in substrate affinity. The doubly truncated Jen1ΔNT94ΔCT33 version and Jen1ΔNT94 also had 2.5 to 3-fold increased affinity for lactate. Thus, transport kinetic parameters for Jen1 truncations revealed that specific segments of the N- and C-termini of the Jen1 transporter are critical for substrate binding and transport dynamics, in addition to their role in PM sorting, stability and regulated endocytosis. These results unmask a previously unnoticed functional interaction of the N- and C-tails, highlighted by the fact that the doubly truncated Jen1 version had an expression and functional profile distinct from the single truncations.

**Figure 3.**
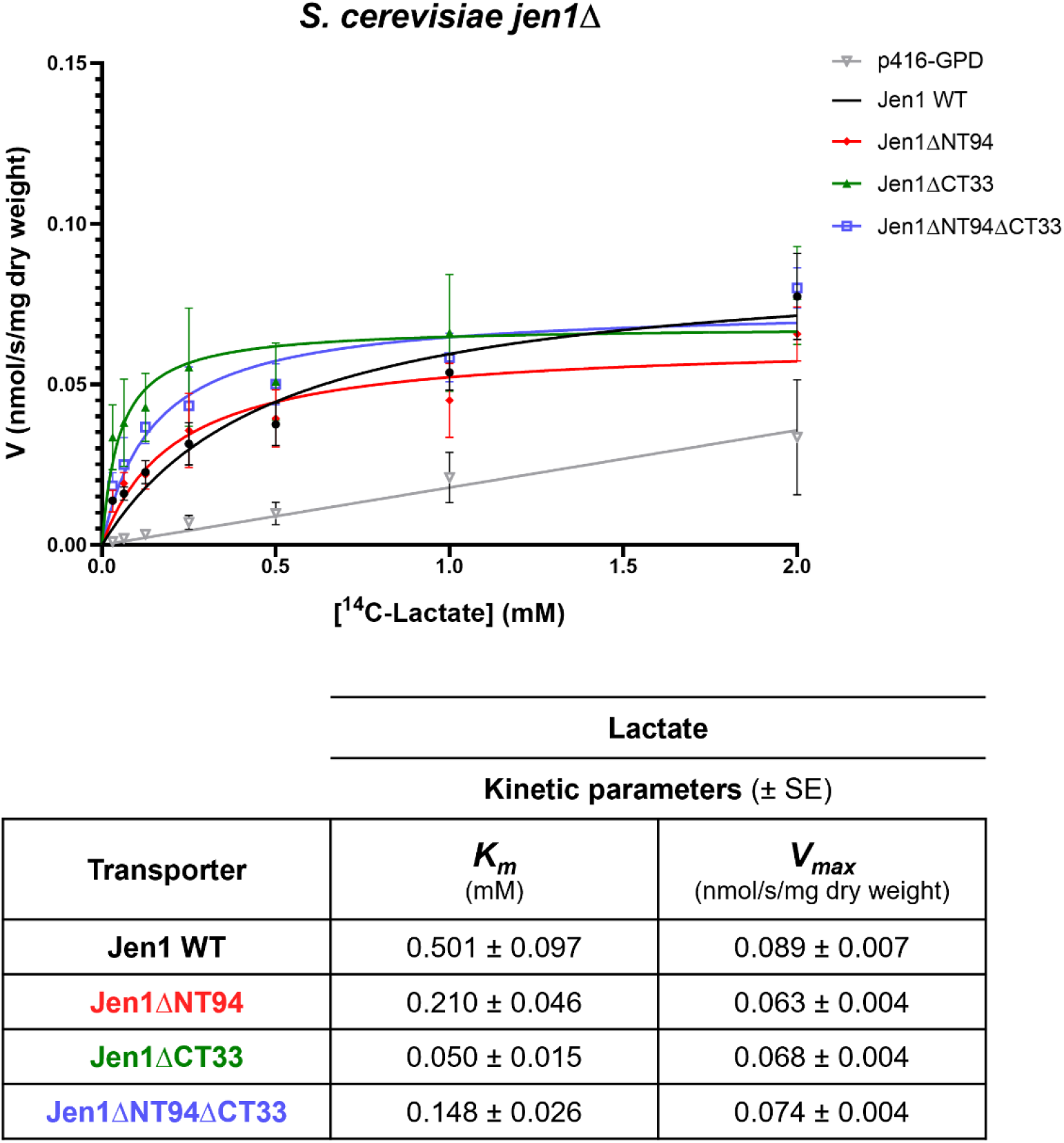
Transport kinetics of Jen1 truncations. The upper panel shows initial uptake rates of radiolabelled ^14^C-lactic acid, pH 5.0, as a function of lactate concentration in *S. cerevisiae jen1*Δ cells expressing *JEN1* gene (Jen1 WT), *JEN1* mutant versions (Jen1ΔNT94, Jen1ΔCT33 and Jen1ΔNT94ΔCT33), or transformed with p416-GPD (empty vector) as a control. The respective kinetic parameters are highlighted in the table (lower panel). The data shown are mean values of at least three independent experiments and the error bars represent the standard deviation. *K_m_* and *V_max_* were determined using the GraphPad Prism 8. *K_m_*, Michaelis-Menten constant; SE, standard error; *V_max_*, maximum velocity.

### Maximal glucose-triggered endocytic turnover of Jen1 involves interactions of Rod1 and Bul1/2 at the C-terminal segment and N-terminus, respectively

The degradation of Jen1 induced by glucose was reported to require both Rod1 and Bul1 arrestins (35, 42). It was thus proposed that multiple α-arrestins may act sequentially to recruit the ubiquitylation machinery preceding endocytosis. Although the specific lysines residues necessary for ubiquitylation of Jen1 still remain under dispute, a glucose-responding degron recognized by Rod1 has been recently identified in the C-terminus of Jen1 (36). No binding motif has been identified for the Bul1 arrestin.

Here, we investigated the localization and protein levels of the Jen1 functional truncations in a standard wild-type background (i.e., *ROD1*+ *BUL1/*2+) and in strains lacking these protein adaptors (i.e., *rod1*Δ, *bul1*Δ*bul2*Δ or *rod1*Δ*bul1*Δ*bul2*Δ) (**Figure 4**, **S3** and **S4**). In these assays, cells were grown under Jen1 induction conditions (Gal 5 h), and then glucose was added for 2 or 4 h to trigger Jen1 internalization (for details see Material and methods). At the times indicated, cells were collected, visualized by fluorescence microscopy and proteins extracts were prepared and analysed by western blot.

**Figure 4.**
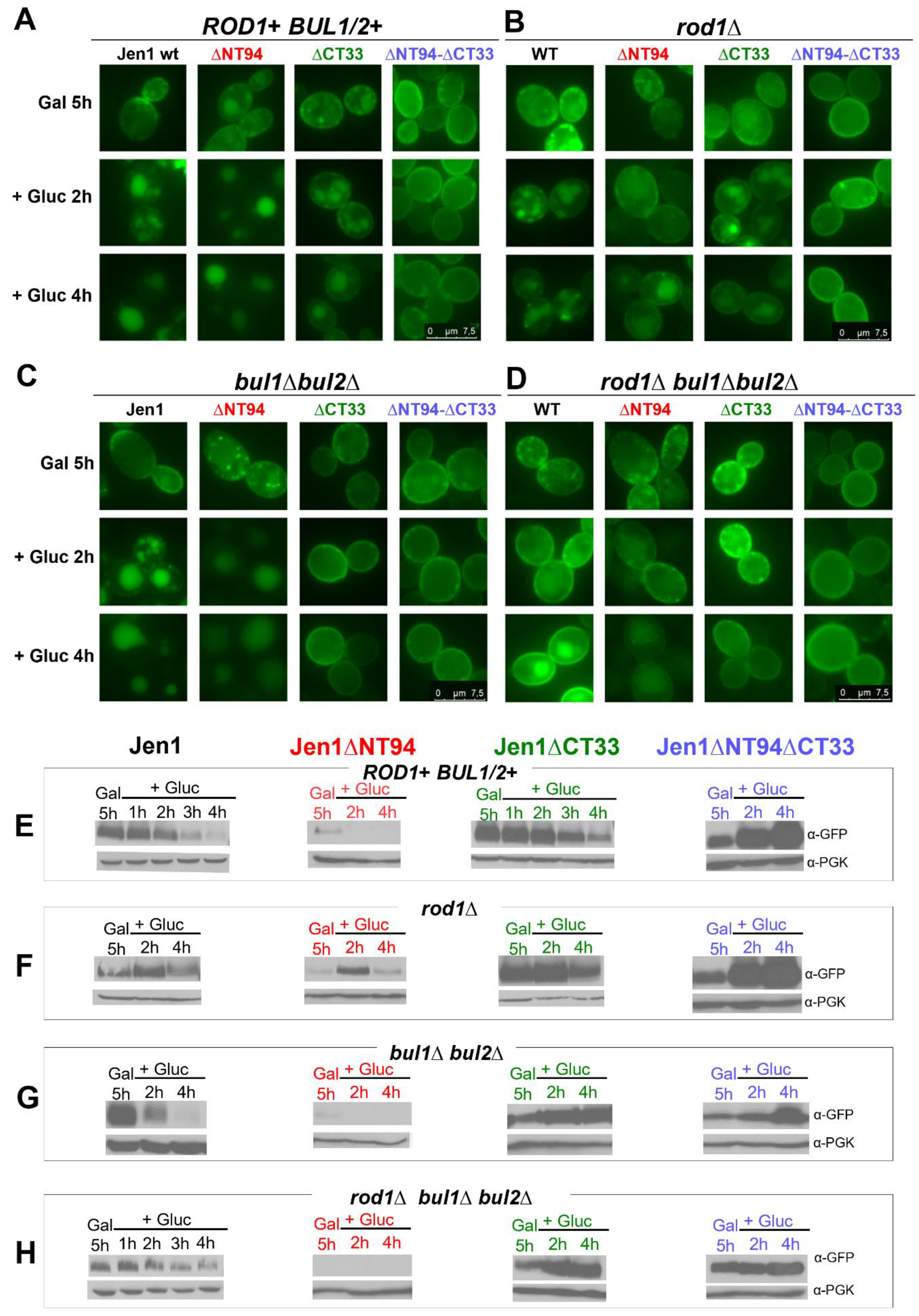
Jen1 N- and C- termini are both involved in glucose-induced down-regulation. Cells of *S. cerevisiae ROD1*+ *BUL1/*2+, *rod1*Δ, *bul1*Δ*bul2*Δ or *rod1*Δ*bul1*Δ*bul2*Δ, expressing the Jen1 truncations tagged with GFP and expressed under a GAL promoter were analysed by epifluorescence microscopy (**A**-**D**) and by western blot (**E**-**H**). Cells expressing Jen1 WT and Jen1 truncations (Jen1ΔNT94, Jen1ΔCT33 or Jen1ΔNT94ΔCT33) were grown overnight in YNB glucose (2 %, w/v) medium and, after being washed twice in deionized water, cells were transferred to YNB galactose (2 %, w/v) medium to induce Jen1 expression of the Jen1 constructs. After 5 h in galactose medium, glucose was added. Cells were visualized by fluorescent microscopy at specific time points (Gal 5 h, Gal 5 h + Gluc 2 h and Gal 5 h + Gluc 4 h). At the same time points, cells were harvested, and protein extracts prepared for Western immunoblotting with an anti-GFP antibody or anti-phosphoglycerate kinase (PGK) antibody (loading control).

Results obtained in the wild-type background showed that wild-type Jen1 is internalized and targeted for vacuolar degradation after glucose addition (**Figure 4A**). This is further confirmed by Jen1 co-localization with CMAC, a blue vacuolar marker (**Figure S3A**), and by the progressive decrease in protein steady state levels (**Figure 4E** and **S4A**). A similar picture in respect to Jen1 localization and stability was obtained in the strain lacking Bul1/2 proteins, suggesting that Bul1/2 participate little, if not at all, in the endocytosis of the wild-type transporter, under our experimental conditions (**Figure 4C** and **4G;** see also **S3C** and **S4**). When expressed in a strain lacking Rod1, the internalization and degradation of Jen1 is still evident, but delayed (**Figure 4B** and **4F**; see also **S3B** and **S4B**). In the triple-deletion mutant, *rod1*Δ*bul1*Δ*bul2*Δ, internalization of Jen1 is very low (**Figure 4D** and **4H;** see also **S3D** and **S4**). Thus, the fact that Jen1 endocytosis is significantly blocked only in the triple mutant indicates a role for both Rod1 and Bul1/2 in glucose-triggered endocytosis, in line with previous reports (35, 42). However, considering the western blots shown in **Figure 4E-H**, the role of Bul1/2 in glucose-triggered endocytosis was found to be more complex, as it seemed to depend on the presence or absence of Rod1. In general, the role of Bul1/2 seemed secondary to that of Rod1 in respect to glucose-elicited endocytosis, as least under our experimental conditions.

Overall, the results obtained concerning wild-type Jen1 point to the conclusion that, in the presence of glucose, Rod1 is the principal arrestin mediating endocytosis, and that Bul1/2 might have a secondary role, especially when Rod1 expression is genetically blocked. To further investigate the possible connection between Jen1 cytosolic tails with the action of Rod1 and Bu1/2 arrestins, we carried out similar experiments using the truncated versions of Jen1.

Fluorescent assays and western blot analysis confirmed that Jen1ΔNT94 is a very unstable Jen1 version, being rapidly degraded after glucose treatment, in the wild-type (**Figure 4A** and **4E;** see also **S4**), but also in the *bul1*Δ*bul2*Δ background (**Figure 4C** and **4G;** see also **S4**). However, Jen1ΔNT94 stability and PM localization increased significantly when *ROD1* was knocked-out (**Figure 4B** and **4F;** see also **S4**). A similar picture was obtained in the triple mutant, *rod1*Δ*bul1*Δ*bul2*Δ (**Figure 4D** and **4H**; see also **S4**). These observations suggest that Rod1 functions via interaction with the Jen1 C-tail, as also previously reported (36). Of note, Jen1ΔNT94 protein levels were extremely low, at the limit of detection, in strains lacking Bul1 and Bul2 proteins (*bul1*Δ*bul2*Δ or *rod1*Δ*bul1*Δ*bul2*Δ) (**Figures 4G** and **4H**).

Fluorescent assays and western blot analysis confirmed that Jen1ΔCT33 is a significantly stabilized Jen1 version, under all conditions tested (**Figure 4A** and **4E**). The absence of Rod1 did not increase further the stability of Jen1ΔCT33 in the PM (**Figure 4B** and **4F**), in line with the fact that Rod1 operates via the ‘missing’ C-tail segment. However, when Bul1/2 are genetically knocked-out, Jen1ΔCT33 was stably localized to the PM, even after 4 h of glucose addition (**Figure 4C** and **4G**). This finding revealed an additive effect of the absence of the last 33 amino acid residues, which include the Rod1 binding site (36), with the absence of Bul1/2 proteins. This result supports the idea that, under glucose-triggered endocytic conditions, Bul1/2 proteins exert their action via the N-tail of Jen1, while Rod1 acts via the C-tail, thus also justifying why Jen1 becomes fully stabilized in the PM when the action of both types of arrestins is genetically suppressed. Notice again that the role of Bul1/2 seems more prominent when Rod1 is missing, suggesting a sequential action of these arrestins. Results obtained in *rod1*Δ*bul1*Δ*bul2*Δ (**Figure 4D** and **4H**) reinforced the aforementioned conclusions. Noticeably also, in the tripe *rod1*Δ*bul1*Δ*bul2*Δ mutant localization of the doubly truncated Jen1 version (Jen1ΔNT94ΔCT33) to the PM is ‘absolute’ and Jen1 protein steady state levels are at their maximum (**Figure 4D** and **4H**), whereas in the same genetic background wild-type Jen1 is undergoing low endocytosis and turnover, best seen after 4 h of glucose addition. This suggests that other arrestins, other than Bul1/2 or Rod1, might still recognize, albeit with lower affinities, the cytosolic tails of Jen1 and thus promote moderate endocytosis. Thus, the truncation of both cytosolic terminal segments of Jen1 proves to be pivotal for generating a Jen1 version that is fully insensitive to endocytosis.

### The C-terminus of Jen1 is sufficient for promoting glucose-elicited turnover of UapA via interaction with Rod1

The UapA transporter heterologously expressed in *S. cerevisiae* is not endocytosed by glucose, ammonium or excess of substrate ((39) and G.D. unpublished). This might be due to the fact that the α-arrestin adaptors of *S. cerevisiae* are not functionally orthologous to those of *A. nidulans* (14) or that the ubiquitylation and/or endocytic machineries in the two fungi are not functionally complementary. We took advantage of the stability of UapA expressed in *S. cerevisiae*, under all conditions tested, to obtain more information on the role of cytosolic terminus domains of Jen1. We thus constructed chimeras between Jen1 and UapA transporters by adding the shorter segments of the cytosolic N- and C-termini of Jen1 to the intact UapA transporter (see **Figure 1B**). For details of strains see Materials and methods. We analysed these chimeras by uptake transport assays, epifluorescence microscopy and western blotting, in different *S. cerevisiae* strains (**Figure 5**).

**Figure 5.**
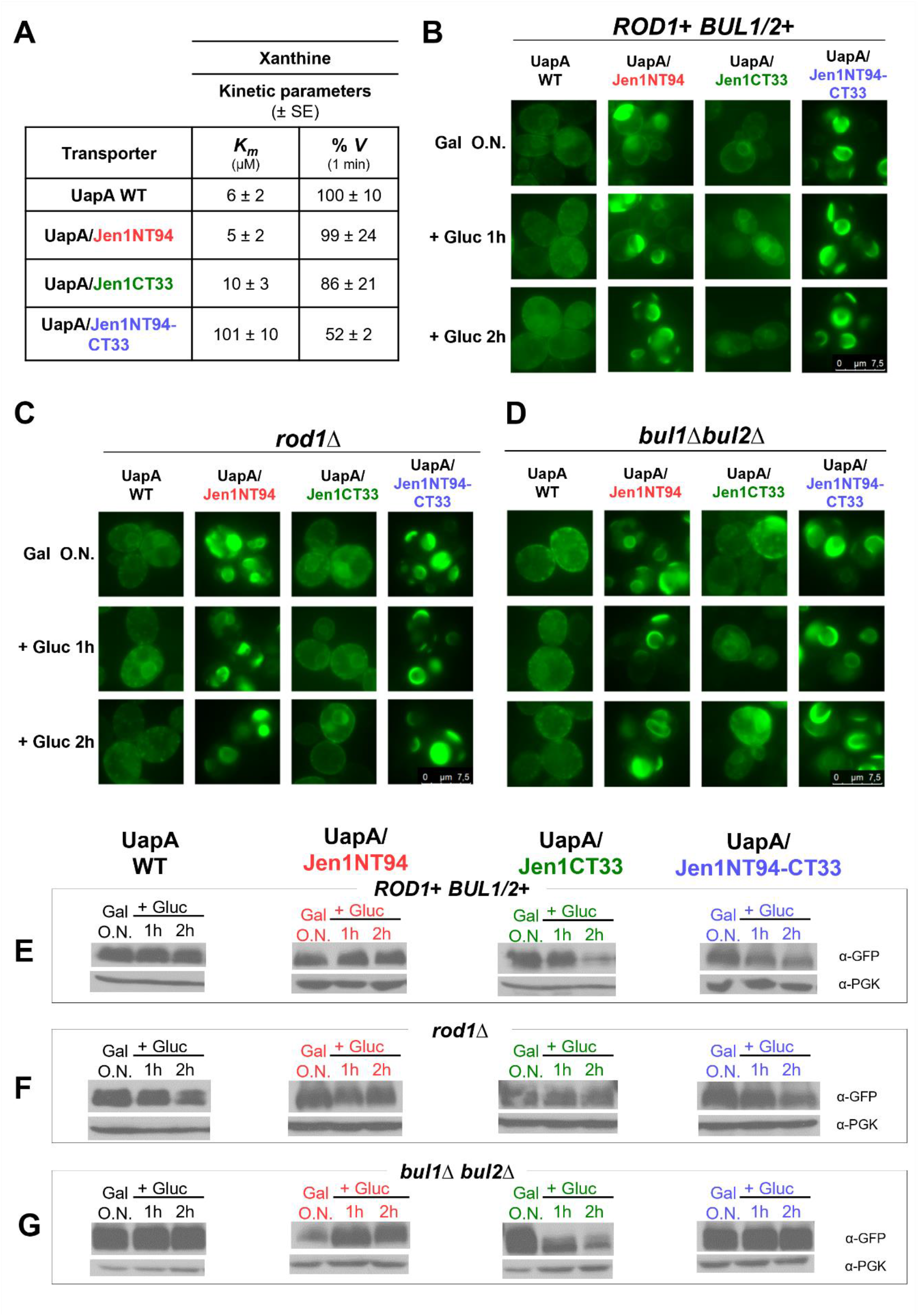
The CT33 segment of Jen1 confers sensitivity to glucose-triggered endocytosis to UapA via interaction with Rod1. Cells of *S. cerevisiae ROD1+ BUL1/2+* strain or cells lacking the arrestins *rod1*Δ or *bul1*Δ*bul2*Δ, expressing UapA-Jen1 chimeras tagged with GFP and expressed under a GAL promoter, were analysed by transport assays using radiolabelled xanthine (**A**), by epifluorescence microscopy (**B**-**D**) and by Western blot (**E**-**G**). Cells were grown overnight (O.N.) in YNB galactose (2 %, w/v) + glucose (0.1 %, w/v) medium until mid-exponential phase and glucose was added when indicated (Gal O.N. + Gluc 1 h and Gal O.N. + Gluc 2 h). At these time points, cells were visualized by Fluorescent Microscopy and protein extracts were prepared for Western immunoblotting with an anti-GFP antibody or anti-phosphoglycerate kinase (PGK) antibody (loading control). Transport assays are described in materials and methods and in (39). *K_m_*, Michaelis-Menten constant; SE, standard error; *V*, relative % velocity.

Transport assays showed that wild-type UapA, UapA/Jen1NT94, UapA/Jen1CT33 and UapA/Jen1NT94-CT33 could all confer saturable xanthine import, showing that a fraction of UapA and the UapA/Jen1 chimeras reach the PM and are transport-active. In particular, UapA and the single chimeras UapA/Jen1NT94 and UapA/Jen1CT33 showed very similar *K*_m_ and *V*_m_ values, similar also to the native *K*_m_ of UapA measured in *A. nidulans* (**Figure 5A**). On the other hand, the ‘double’ chimera UapA/Jen1NT94-CT33 showed reduced transport function, as the relevant *K*_m_ and *V*_m_ values were increased and reduced, respectively (**Figure 5A**), an indication that this chimera might be partially misfolded when expressed in yeast.

The localization of wild-type UapA and the three chimeras (UapA/Jen1NT94, UapA/Jen1CT33 and UapA/Jen1NT94-CT33) was followed in a standard wild-type *S. cerevisiae* carrying a *jen1* deletion (*jen1*Δ) and in a strain lacking all six ER-PM tethering proteins. **Figure 5B** (upper row) shows that in a standard wild-type background, upon transcriptional induction (Gal O.N.), UapA and the ‘single’ chimeras UapA/Jen1NT94 and UapA/Jen1CT33, all showed significant PM localization, concomitant however with partial retention in the ER, especially in the case of UapA/Jen1NT94, as revealed by the fluorescent labelling of perinuclear ER membranes. The double chimera UapA/Jen1NT94-CT33 seemed to be massively retained in ER-like cytosolic structures (**Figure 5B**). When expressed in the strain lacking the ER-PM tethering proteins (Δtether), UapA and all chimeras tested showed increased, but variable, PM localization (**Figure S6**). More specifically, UapA labeled exclusively the PM, UapA/Jen1CT33 showed strong PM labelling and very minor ER retention, UapA/Jen1NT94 localized mostly in the PM and partial retention in perinuclear ER rings, while the great majority of UapA/Jen1NT94-CT33 molecules were retained in the ER. These results confirmed those obtained in the standard genetic background used in most of this study. Thus, our findings showed that all UapA-Jen1 chimeras translocate to the PM, albeit with different efficiency, which was in accordance with uptake assays showing that chimeras are functional, as they all mediate xanthine import (**Figure 5B** and **S5**). Noticeably, expression of UapA and UapA-Jen1 chimeras in the Δtether strain showed significantly increased translocation to the PM, compared to the expression in our standard yeast strain. In other words, abolishment of cER-PM contacts enhanced translocation of UapA and chimeras to the PM. This is a rather surprising and interesting result that needs to be followed in the future, given it falls beyond the scope of the present work.

After having established that UapA and UapA-Jen1 chimeras are functionally translocated to the yeast PM, we followed their response to glucose-triggered endocytosis. **Figure 5B** (middle and lower rows) shows that in the presence of glucose the localization profile of UapA, UapA/Jen1NT94 and UapA/Jen1NT94-CT33 was similar with that obtained without glucose, suggesting that the relevant proteins are insensitive to glucose-triggered endocytosis. In contrast, UapA/Jen1CT33 showed some evidence for glucose triggered endocytosis, clearer after 2 h of glucose addition. This conclusion was well supported by the observation that UapA/Jen1CT33 co-localized with the blue vacuolar marker, a result not observed with UapA or the other chimeras (**Figure S5A**). To further confirm the response of UapA/Jen1CT33 to glucose-elicited endocytosis, we measured the steady state protein levels of these proteins by western blotting. As shown in **Figure 5E**, solely UapA/Jen1CT33 protein levels were significantly reduced in the presence of glucose. This suggests that the C-terminus of Jen1 is sufficient to promote glucose-elicited turnover of UapA, in a context-independent manner.

We, subsequently, analysed the localization and the protein steady state levels of UapA and UapA-Jen1 chimeras in *rod1*Δ and *bul1*Δ*bul2*Δ strains, in the absence or presence of glucose. In both strains and in all conditions tested, wild-type UapA was stably translocated to the PM with some evidence of moderate ER-retention (**Figure 5C**, **5D**, **5E**, **5F**, **5G** and **S5A**). This confirmed that wild-type UapA does not respond to endocytosis in yeast, and very probably it is not recognized by Rod1 or Bul1/2. UapA/Jen1CT33, a chimera shown previously to respond to endocytosis by glucose in a wild-type background (i.e., *ROD1+BUL1/2+),* when expressed in *rod1*Δ strain remained stably localized to the PM irrespective of presence or absence of glucose (**Figure 5C**, **5F** and **S5B**). This result, best highlighted when comparing western blots in **Figure 5E** and **5F**, showed that UapA/Jen1CT33 endocytosis is mediated by Rod1, in line with the idea that Rod1 binds to the C-terminal segment of Jen1. UapA/Jen1CT33 biogenesis in *bul1*Δ*bul2*Δ was less clear. In the microscopy analysis, no fluorescence could be detected, as probably expected, a convincing response to glucose-triggered endocytosis (**Figure 5D** and **S5C),** but in western blots it was clear that the steady state levels of this chimera are down-regulated by glucose, similar to the result obtained in the *ROD1*+*Bul1/2*+ wild-type background (**Figure 5G**). Thus, Bul1/2, unlike Rod1, did not seem to contribute to UapA/Jen1CT33 endocytosis.

Subcellular localization results, obtained with UapA/Jen1NT94 and especially with UapA/Jen1NT94-CT33, were more complex to interpret regarding the role of Jen1 terminal regions, mostly due to significant ER-retention of these chimeras (**Figure 5C, 5D, S5B** and **S5C**). Based solely on the western blot analysis (**Figure 5E, 5F** and **5G**), we may conclude that none of the two chimeras responds to glucose-triggered endocytosis, given that their steady state levels were little changed both in the presence and in the absence of glucose.

Overall, our results supported the concept that Rod1 binds directly to the Jen1 C-terminal 33 amino acid residues added in the tail of UapA, and thus promotes endocytosis of the respective chimera, in response to glucose. This further shows that interaction of Rod1 with a specific sequence motif present at the C-terminal of Jen1 is context-independent. On the other hand, the presence of the Jen1 N-terminus did not seem critical for the turnover of UapA, suggesting that the proposed interaction of Bul1/2 proteins with this region might be weaker and/or context-dependent.

### Dynamic transport-dependent interaction of N- and C-termini of Jen1

Overall, our genetic and biochemical results strongly suggested that N- and C-termini of Jen1 contain elements critical for biogenesis, function and turnover of Jen1. Most notably, the effects of Jen1 cytosolic termini on Jen1 functional expression proved additive. This was highlighted by the significant stabilization of Jen1, when truncated at both tails, in comparison to what was found for the singly truncated versions. These results also suggest that the termini of Jen1 might interact with each other during the conformational changes accompanying transport activity, which is also associated to endocytic turnover. To further investigate this issue, we used a Bimolecular fluorescence (BiFC) assay, based on reconstitution of YFP florescence when the two parts of the split YFP epitope are fused in the two tails of Jen1 (YFPn-Jen1-YFPc). Given that reconstitution of YFP might in principle also occur in case Jen1 dimerizes, we also constructed a strain co-expressing the two parts of the split YFP epitope fused in distinct Jen1 molecules (i.e., Jen1-YFPn or Jen1-YFPc). These strains and relative controls (i.e., strains expressing Jen1-YFPn or Jen1-YFPc or co-expressing both) were used to investigate whether YFP is reconstituted in *cis* via interaction of the Jen1 tails, or/and in *trans* via dimerization of Jen1 molecules (for details of constructs see material and methods). **Figure 6A** shows that strong, PM-associated, reconstitution of YFP fluorescence occurs solely when the split epitope parts are fused with the tails of Jen1, whereas no fluorescent signal was obtained when these are fused in different Jen1 molecules. This result not only strongly suggests that the two tails of Jen1 come in close contact when attached in the same Jen1 molecule, but also points against tight dimerization of distinct Jen1 molecules, at least under the conditions tested. To address further the mechanism by which the two tails of Jen1 come into contact, we repeated our assay in the presence of substrate. **Figure 6B** shows that the presence of lactic acid reduced significantly the YFP fluorescence signal over time (most evident at 30 min), while after prolonged incubation this was stabilized at a very low basal level (50 min). This finding suggests that stable reconstitution of YFP is transport-activity dependent, as the conformational movements accompanying translocation of the substrate should in principle affect the positioning of the two tails in the outward- and inward-facing topologies of Jen1. The transport-dependent interaction of N- and C-termini of Jen1 is very similar to what has been observed in members of an evolutionary, structurally, and functionally distinct transporter family, namely the NCS1/APC superfamily (33).

**Figure 6.**
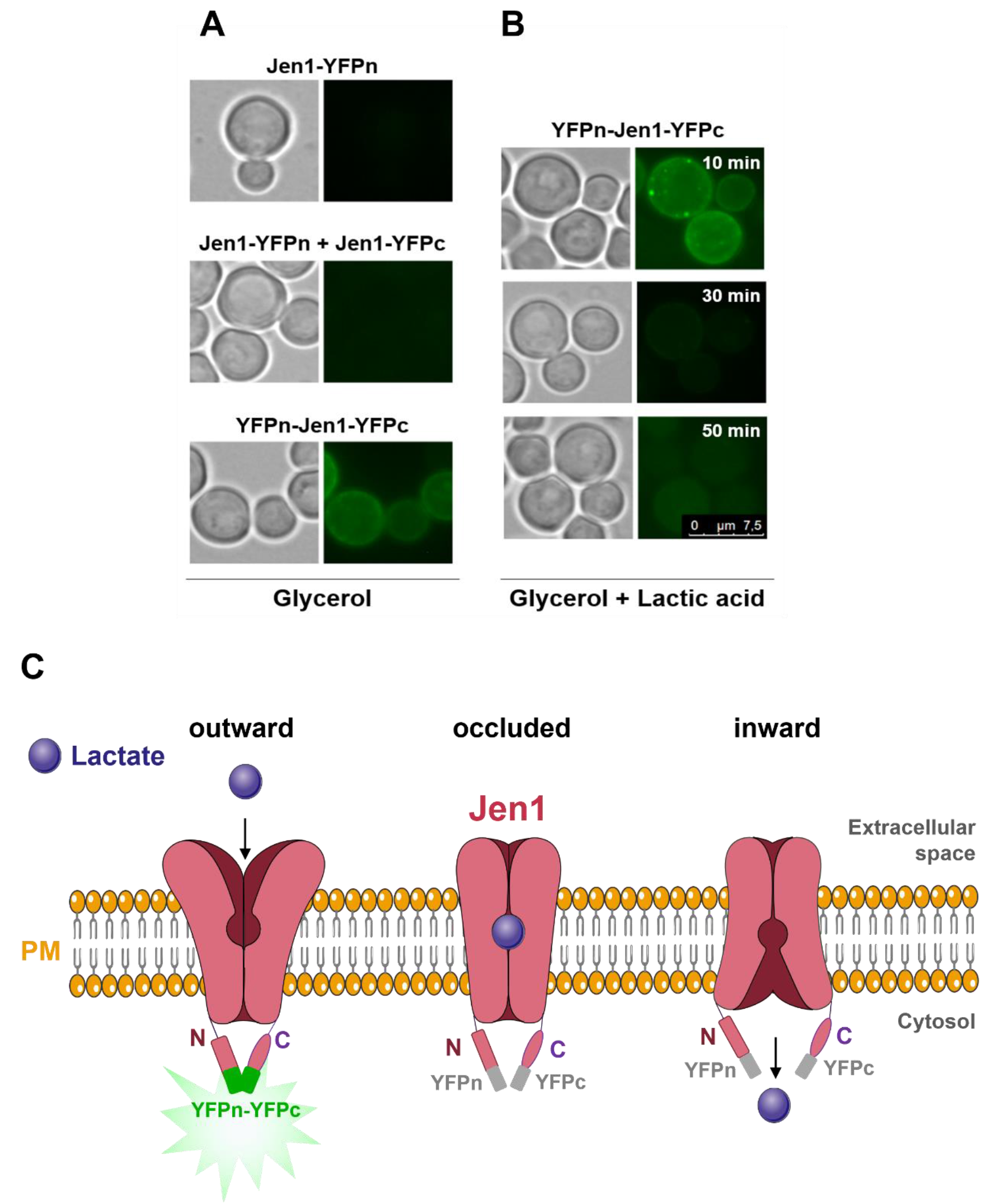
The cytosolic termini of Jen1 come into close proximity in the absence of its substrate. Cells of *S. cerevisiae jen1*Δ strain expressing YFPn-Jen1-YFPc, Jen1-YFPn, or co-expressing both Jen1-YFPn and Jen1-YFPc were analyzed by *in vivo* epifluorescence microscopy (for more details see Materials and Methods). (**A**) To induce Jen1 expression at the plasma membrane (PM), cells were grown in YNB 3 % (v/v) glycerol, until mid-exponential phase. (**B**) 0.5 % (v/v) lactic acid was added to cells previously grown in YNB 3 % (v/v) glycerol, until mid-exponential phase, and samples were collected over time for fluorescent microscopy analysis. (**C**) Schematic representation of the Jen1 transporter termini conformation during a substrate transport cycle (i.e., outward-facing, occluded, inward-facing). In the absence of substrate, Jen1 is in an outward-facing conformation with the N- and C-termini in close contact or interacting with each other. Upon substrate addition, Jen1 moves to an inward conformation with consequent loss of termini interaction/proximity. The topological changes in Jen1 termini seem to be crucial for Jen1 endocytic turnover, biogenesis/folding, transport activity and trafficking. The aforementioned scheme represents a speculative *model* supported by our results.

## Discussion

In the present work, we investigated possible functional roles of the terminal cytosolic segments of the Jen1 monocarboxylate transporter in *S. cerevisiae* (see **Table S2**). Our first approach consisted in designing, genetically constructing and functionally analysing truncated versions of GFP-tagged Jen1, lacking parts of their cytosolic termini, expressed in wild-type or mutant *S. cerevisiae* strains lacking the arrestins Rod1 or Bul1/2. Our functional analyses included Jen1-mediated growth tests on lactic acid, or effect on external pH, direct measurements of Jen1 transport kinetics using radiolabelled lactic acid, *in vivo* imaging of subcellular localization, and western blot measurements of protein steady state levels of the truncated Jen1 versions. These constructs were expressed under induction conditions and in response to physiological signals triggering Jen1 endocytosis (i.e., glucose or prolonged growth on lactate).

Subsequently, we generated and analysed functional chimeric transporters made of UapA, a heterologous nucleobase-allantoin transporter of *A. nidulans,* fused with the terminal regions of Jen1.

Deleting the entire Jen1 terminal regions, which correspond to 133 N-terminal or 62 C-terminal amino acids, as defined by *in silico* predictions, led to non-functional Jen1 versions, which in most cases were associated with significant cellular mislocalization, mostly ER-retention. We thus proceeded by analysing shorter truncations, as those deleting the 94 N-terminal or/and the 33 C-terminal amino acids. Notice that similar truncated versions of Jen1 have been previously analysed (36), which allowed direct comparison of relevant results (see later). The shorter Jen1 truncations were proved to be functional based on growth tests and other functional assays, similarly to what has been reported in (36). Based on these Jen1 truncations and relative chimeras with UapA, we came to the following primary observations.

### Role of the C-tail

Jen1ΔCT33 is a stable version of Jen1, in all conditions tested, showing also increased *capacity* of lactate transport not only because of its higher concentration in the PM, but also due to 10-fold increased affinity for its substrate. This suggests that the segment of the C-terminal 33 amino acids deleted in the truncated version, contains not only a degron, as reported by Fujita et al (2018) (36), but also a functional motif that seems to affect the mechanism of transport of Jen1 in an ‘allosteric’ manner. This conclusion is based on the fact that although the structure of Jen1 is not known, but only predicted via homology modeling, its cytosolic C-tail is, in principle, distant from the proposed substrate translocation trajectory (43). A similar situation of regulation of transport kinetics by genetic modifications of the cytosolic C-tail has been reported for FurE, an *A. nidulans* nucleobase-allantoin transporter (32, 33). We also present evidence that the C-terminal 33 amino acid segment of Jen1 contains the major Rod1 interacting motif, as also reported in Fujita et al. (2018) (36), given that its deletion (i.e., ΔCT33) mimics the absence of glucose-triggered endocytosis of Jen1, observed in a *rod1* null mutant. Additionally, in the absence of Bul1/2, full endocytosis is observed in wild-type Jen1 and Jen1ΔNT94, but not in Jen1ΔCT33. Furthermore, we show that the interaction of Rod1 with the 33 amino acid C-terminal segment of Jen1 is *direct* and *context-independent*, because its transfer to the endocytosis-insensitive UapA proved sufficient to promote Rod1-dependent down-regulation, in the presence of glucose. Overall, our results concerning the C-terminal part of Jen1 confirm the conclusions presented in Fujita et al (2018), but further reveal two novel properties of this part of the transporter. First, the C-tail of Jen1 regulates the transport mechanism from a distance, and second, Rod1 recognizes a motif in the C-tail of Jen1 without the involvement of other regions of the transporter. To our knowledge there is no other report showing a context-independent interaction of a transporter motif with α-arrestins.

When considering the fact that Jen1ΔCT33 is stable and fully functional while Jen1ΔCT62 is non-functional due to retention in the ER, we can also conclude that the middle part of the C-terminal segment, between amino acids residues 554-583 (the areas missing in Jen1ΔCT62 but present in Jen1ΔCT33, **Figure S2**), might contain elements critical for ER-exit or proper folding. Such ER-exit motifs have been identified at the cytosolic termini of other transporters but, to our knowledge, none has been rigorously shown to act in a context-independent manner (see the recent review (34). A preliminary *in silico* analysis of the sequence of Jen1 between amino acids residues 554-583 showed that a short di-acidic motif, ^577^EYE^579^ might be an interesting candidate as an ER-exit motif (see **Figure S2B**).

### Role of the N-tail

Jen1ΔNT94 was shown to be normally produced at basal levels, but proved to be a rather unstable version of Jen1, exhibiting rapid turnover upon further induction. Thus, the N-terminal 94 residue segment of Jen1 should include elements critical for post-translational stability, evident upon translocation to the PM. Interestingly, Jen1ΔNT94 showed moderately altered transport kinetics (e.g., 2.5-fold increased substrate affinity), which points to a positive ‘distant’ effect on the transport mechanism, albeit weaker than that of the C-terminal segment. Notably also, the N-terminal part proved critical for endocytic down-regulation in response to prolonged growth on lactate or glucose, because when *ROD1* gene was knocked-out or the C-tail of Jen1 was deleted (i.e., no Rod1 binding), the presence of the N-terminal conferred partial endocytosis, while its absence led to an increased stability. Our data further support the conclusion that glucose triggered endocytosis of Jen1 is exerted via Bul1/2 binding to the N-terminal segment, as endocytosis without the C-terminal region (Jen1ΔCT33) or without an active *ROD1*, depends solely on Bul1/2.

When the NT94 segment of Jen1 was transferred to UapA, it led to a chimera that showed significant ER-retention, despite being transport-competent. As a result, the functional analysis of this chimera did not provide us with additional evidence on the role the Jen1 N-tail. Finally, comparing the effect of deleting the entire N-terminal cytosolic region of Jen1 (residues 1-133), which led to ER-retention, to Jen1ΔNT94, which led to turnover after translocation to the PM, we conclude that the segment between 94-133 might also include motifs driving ER-exit or necessary for proper folding. Interestingly, this Jen1 segment contains a well-conserved motif, ^126^NPIPE^133^, that is worthy to be studied by mutational analysis, in the future (see **Figure S2B**).

### Dynamic interactions of the N- and C-tails of Jen1

A clear conclusion concerning the tails of Jen1 is that both are needed for maximal glucose triggered endocytosis of Jen1, with the N-tail interacting with Bul1/2 and the C-tail with Rod1. The interaction of Rod1 with the C-tail seems to result in a stronger Jen1 endocytosis when Bul1 interaction with the N-tail is blocked, while the interaction of Bul1/2 with the N-tail confers only partial endocytosis, when Rod1 interaction with the C-tail is genetically abolished. To our opinion, however, the most interesting novel finding of this work concerns the evidence supporting the conclusion that the two Jen1 termini co-operate in regulating the stability and function of Jen1. A first genetic indication supporting this idea came from the doubly truncated Jen1 version, which showed exceptional new properties, other than those of the singly truncated mutants. More specifically, Jen1ΔCT33ΔNT94 shows very high PM stability, under all conditions and genetic backgrounds tested, displaying a transport kinetics distinct from the single truncations and the wild-type Jen1. Thus, the doubly truncated Jen1 version is a ‘new’ monocarboxylate transporter that is endocytosis ‘resistant’ and that has an increased substrate affinity relative to the wild-type Jen1.

To address the molecular basis underlying the additive functional roles of Jen1 tails, we employed a BiFC assay, which showed a dynamic and transport-dependent interaction of the two tails of Jen1. Using the same assay, we also obtained strong evidence that Jen1 does not form dimers, at least in the conditions tested, which proved fortuitous for more rigorously interpreting the positive BiFC signals obtained when the two parts of split YFP were cloned in the same Jen1 molecule. The only other previously reported case where BiFC assays showed that cytosolic tails interact to control the function and turnover of a transporter, is that of FurE in *A. nidulans* (32, 33). In this case, interactions of the two tails affected the stability, trafficking, function and endocytosis of FurE, and surprisingly, substrate specificity. Preliminary Molecular Dynamic analysis has provided some hints on how cytosolic tails might have affected FurE functioning from a distance by modifying the opening and closing of outer and inner gates of the transporter (33). In the present work, Jen1 truncations did not seem to affect substrate specificity, but interestingly, all functional Jen1 truncations showed increased (2.5 to 10.0-fold) affinities for lactic acid transport, despite retaining wild-type *V*_m_ values (see **Figure 3**). The alteration in *K*_m_ values reveals a modification in the capacity of Jen1 truncations to bind native substrates. In other words, similar to FurE, changes on the cytosolic tails of Jen1, distantly positioned from the substrate binding site, affect the mechanism of substrate selection and transport. How this is achieved in the case of Jen1 remains elusive, but constitutes an interesting point to be addressed in the future.

Finally, another interesting observation of this work concerns the finding that UapA-Jen1 chimeras are more massively translocated into the PM when cER-PM contacts are abolished. The molecular basis of this phenomenon is still unclear. In fact, we did not notice a general enhancement of sorting to the PM of wild-type Jen1 when cER-PM contacts are abolished. It thus seems that vesicular trafficking of specific cargoes, such as the UapA-Jen1 chimeras, is enhanced when the cER is not in close proximity to the PM. This finding might prove extremely valuable for the expression of heterologous membrane proteins in yeast.

The present work on Jen1 also shows that rather cryptic roles of transporter cytosolic tails can be exploited to rationally modify transporter function, which will be valuable, not only for addressing basic mechanisms of solute transport, but also for serving as tools in biotechnological applications ((44), for a review see (8)). The generality of this concept is supported by the work on FurE and Jen1, representing the two major transport superfamilies, APC and MFS, but also several other reports directly or indirectly supporting the emergence of transporter tails as important functional elements (34).

## Supporting Information

This article contains supporting information.

## Materials and Methods

### Yeast strains and Growth conditions

All the yeast strains used in this work are listed in **Table S3**. The strains *jen1*Δ, *rod1*Δ, *bul1*Δ*bul2*Δ and *rod1*Δ*bul1*Δ*bul2*Δ were derived from the 23344c wild type strain (Laboratory collection). For BiFC analysis, a *jen1*Δ strain derived from W303-1A was used (Laboratory collection). Yeast cells were grown in a synthetic minimal medium with 0.67 % (w/v) yeast nitrogen base (Difco), supplemented to meet the auxotrophic requirements (YNB medium) or in yeast extract (1 %, w/v) and peptone (1 %, w/v) (YP medium). Solid media was prepared adding agar (2 % w/v) to the respective liquid media. Carbon sources utilized were glucose (2 %, w/v), lactic acid (0.5 %, v/v, pH 5.0), galactose (2 %, w/v) or glycerol (3 %, v/v). Growth was carried out at 30 °C. Cultures were harvested during the mid-log phase of growth. Glucose-grown cells were, then, centrifuged, washed twice in deionized water and cultivated into a fresh YNB medium with lactic acid (incubation time is indicated). For induction conditions of the GAL promoter, the YNB medium was supplemented with a complete mixture Drop-out–uracil + 40 adenine (Formedium). Cells were grown overnight (till an OD_640_ of 1.2-1.8) in YNB medium with 2 % (w/v) glucose and then, after being washed twice in deionized water, they were transferred to YNB medium with 2 % (w/v) galactose at a starting OD_640_ of 0.2. Alternatively, cells were grown overnight (till an OD_640_ of 0.5) in YNB medium with 2 % (w/v) galactose (plus 0.1 %, w/v, glucose), as described by (39). Glucose (2 %, w/v) was added, when indicated.

### Bioinformatic tools

The protein sequences were obtained in SGD (http://www.yeastgenome.org) and AspGD (http://www.aspgd.org/) databases. The secondary structures were predicted by TOPCONS (45). Tertiary structures were predicted by HHpred (http://toolkit.tuebingen.mpg.de/hhpred) and MODELLER software (46), as described previously (43, 47). The minimum number of residues predicted in this work for N- or C-terminus of Jen1 are listed in **Table S1**.

### Construction of transporter truncations and chimeras

All constructions were performed by *in vivo* gap repair (48). Firstly, DNA fragments were amplified by PCR (Accuzyme DNA Polymerase, Bioline, or Supreme NZYProof DNA Polymerase, Nzytech) with specific oligonucleotides (listed in **Table S4**) using yeast genomic DNA (unless it is clearly specified). The resulting PCR products were co-transformed with a linearized plasmid (digested with a specific restriction enzyme) in *S. cerevisiae* cells. All plasmids used and constructed are listed in **Table S5**. Specifically, for construction of *JEN1* termini truncated versions pGPDJEN1ΔNT133, pGPDJEN1ΔCT33, pGPDJEN1ΔCT62, *JEN1* gene DNA fragments were amplified using the following oligonucleotides pairs, respectively: D-NTJEN1_133 and RCTJEN1; D-CTJEN1_33 and CYC1TERM; and D-CTJEN1_62 and CYC1TERM. The resulting PCR products were co-transformed with the linearized plasmid pGPDJEN1. For pGPDJEN1ΔNT94 and pGPDJEN1ΔNT94ΔCT33 constructions, the DNA fragments were amplified from pGPDJEN1 and pGPDJEN1ΔCT33, respectively, using the oligonucleotides Fw_GPD_jen1dNT94 and Rev_GFP_jen1. The resulting PCR products were co-transformed with the linearized plasmid p416GPD. For construction of *JEN1* termini truncated versions under the control of the GAL promoter: pGALJEN1ΔCT33, pGALJEN1ΔNT94 and pGPDJEN1ΔNT94ΔCT33, the *GAL* DNA fragment was amplified from pGALJEN1 with the oligonucleotides GPDfwd and GALrev for pGALJEN1ΔCT33 construction or with the oligonucleotides GPDfw_new and GALrev_dNT94 for constructions pGALJEN1ΔNT94 and pGPDJEN1ΔNT94ΔCT33. The resulting *GAL* PCR product were co-transformed with the respective linearized plasmid (pGPDJEN1ΔCT33, pGPDJEN1ΔNT94 or pGPDJEN1ΔNT94ΔCT33). For the construction of pGALUAPA/JEN1CT33, the *GALUAPA* DNA fragment was amplified from pDDGFP2UAPA using the primers 381 and UapA_rev_33; the *JEN1CT33* DNA fragment was amplified from pDDGFP2UAPAΔCT/JEN1CT62 using the primers Jen1_fw_ct33 and 317. These DNA fragments were then co-transformed with linearized p426GPD plasmid previously digested with *SacI* and *XhoI* restriction enzymes to remove the GPD promoter. For the constructions of pGALUAPA/JEN1NT94, the *GALJEN1NT94* DNA fragment was amplified from pGALJEN1 using the primers 381 and REV_Jen1_UapA; the *UAPAGFP* DNA fragment was amplified from pDDGFP2UAPA using the primers FW_UapA_Jen1 and 317. These DNA fragments were then co-transformed with linearized p426GPD plasmid previously digested with *SacI* and *XhoI* restriction enzymes to remove the GPD promoter. For the constructions of pGALUAPA/JEN1NT94-CT33, *GALJEN1NT94* DNA fragment was amplified from pGALJEN1 using the primers 381 and REV_Jen1_UapA; the *UAPA/JEN1CT33GFP* DNA fragment was amplified from pGALUAPA/JEN1CT33 using the primers FW_UapA_Jen1 and 317. These DNA fragments were then co-transformed with linearized p426GPD plasmid previously digested with *SacI* and *XhoI* restriction enzymes to remove the GPD promoter. The pDDGFP2JEN1ΔCTJEN1CT62 plasmid was derived from pDDGFP2UAPA (Leung et al., 2010). Plasmid isolation from *S. cerevisiae* and *E. coli* strains was performed by standard protocols. Transformations were performed by the standard lithium acetate/polyethylene glycol method (49). All constructs were confirmed by DNA sequencing (GATC Biotech and MWG Eurofins).

### Transport assays

Transport activity assays for Jen1 WT and Jen1 truncated transporters were performed as previously described (47) using radiolabelled D,L-[^14^C] lactic acid (Amersham Biosciences). Yeast cells were mid-exponentially grown in glucose and transferred to a fresh 0.5 % (v/v) lactate medium for 4h (Lac 4h). For uptake measurements, yeast cells were harvested in Lac 4h and centrifuged (5000 rpm, 2 minutes). The samples were then washed twice with ice-cold deionized water and resuspended in ice-cold deionized water to a final concentration of about 20–40 mg dry weight/mL. The reaction mixtures were prepared in 1.5 mL tubes containing 60 μL of KH_2_PO_4_ (0.1 M, pH 5.0), and 30 μL of the yeast cell suspension. After incubation, the reaction was started by the addition of 10 μL of 6 mM radiolabelled lactic acid, pH 5.0 0 (specific activity 2000 dpm/nmol), rapidly mixed by vortexing, and incubated for 1 min. After one minute, 100 μL of 100 mM non-labelled substrate, was added, quickly mixed by vortexing and the mixture was chilled on ice, to stop the reaction. The reaction solutions were centrifuged for 5 min at 13200 rpm. The supernatant was rejected, and the pellet was resuspended in 1 mL of deionized cold water and centrifuged for 5 min, at 13200 rpm. The resulting pellet was resuspended in 1 mL of scintillation liquid (Opti-Phase HiSafe II; LKB FSA Laboratory Supplies). Radioactivity was measured in a Packard TRI-CARB 4810 TR liquid scintillation spectrophotometer with disintegrations per minute correction. The % uptake rate of wild-type Jen1 (Jen1 WT) is considered 100 %. The data is represented as a scatter plot with bar (mean and SD) (GraphPad Prism 8 software) of all data points obtained in three independent experiments. For kinetic assays the methodology used was the same as described above for transport activity assays. However, in this case, the cells were exposed for 30 seconds to various concentrations of radiolabelled D, L-[^14^C] lactic acid, ranging from 0.03 to 2 mM. The data is represented as a Michaelis-Menten plot of the net initial velocity relative to increasing lactic acid concentrations, showing the mean values of at least three independent experiments. The error bars represent the standard deviation. *K_m_* and *V_max_* were determined using the GraphPad Prism 8. Transport activity assays for wild-type UapA and UapA-Jen1 chimeras were performed essentially as described in Leung et al., 2010, using radiolabelled [^3^H]-xanthine (21.1 Ci/mmol, Moravek Biochemicals, Brea, CA).

### Phenotypic growth tests

Phenotypic growth assays on solid medium were performed according to (16, 43). A serial of 1:10 yeast cell dilutions (starting from an OD_640_ of 1) were performed and 3 μL of each yeast suspension were plated in YNB solid medium, containing the desired carbon source. Cells were incubated at 30 °C or 18°C, for 4 or 7 days, respectively.

### Epifluorescence microscopy

Yeast cells were grown, as described above, and visualized by fluorescent microscopy. A volume of 1 mL of growing yeast cells was collected and concentrated by a factor of 10. 5 μL of each sample was then directly visualized, without fixation, on a Leica DM5000B microscope with appropriate filters. The resulting images were acquired with a Leica DFC 350FX R2 digital camera using the LAS AF software. Images were then processed in the Adobe Photoshop CC 2018 (Adobe Systems).

### Measurement of yeast culture pH

The pH of the culture medium was determined as previously described (16). A volume of 1 mL of cell culture was harvested and the pH value was immediately measured by a pHmeter (Braun). The data is represented as an interleaved scatter plot (mean and SD) (GraphPad Prism 8 software) of at least three independent experiments.

### Western blot analysis

Yeast cells were grown as above-mentioned and crude protein extracts were prepared as previously described (50). Nitrocellulose membranes (GE Healthcare Life Sciences) were probed with anti-GFP (clones 7.1 and 13.1, Roche) and anti-PGK (yeast 3-phosphoglycerate kinase, Invitrogen) antibodies, used at 1:3000 and 1:10000 dilutions, respectively. Primary antibodies were detected with horseradish peroxidase-conjugated anti-mouse immunoglobulin G (Sigma) by enhanced chemiluminescence.

### For bimolecular fluorescence complementation (BiFC) assay

For BiFC analyses, several plasmid constructions were performed (**Table S4**): pGPDJEN1YFP_N_ (URA3), pGPDJEN1YFP_C_ (HIS3), pGPDYFP_N_JEN1YFP_C_ (URA3), pGPDYFP_N_ (URA3) and pGPDYFP_C_ (URA3), using a GAP repair cloning strategy. The N-terminal half of the yellow fluorescent protein (YFP_N_; 154 AA residues of YFP), and the C-terminal half of YFP (YFP_C_; 86 AA residues of YFP) were amplified from plasmids PDV7 and PDV8 (51), respectively. *JEN1* ORF was amplified from pGPDJEN1 plasmid. The primers used are listed in **Table S3.**

To study the possible interaction of the Jen1 N- with C-terminus, *jen1*Δ cells expressing pGPDYFP_N_JEN1YFP_C_ were grown overnight in glycerol (3 %, v/v) and ethanol (1 %, v/v), supplemented with the required auxotrophies, until mid-exponential phase, to induce Jen1 expression at the PM. Then, a pulse of lactate (0.5 %, v/v) was added, and fluorescent images were acquired at the indicated time points. Cells co-expressing pGPDJEN1YFP_N_ (URA3) and pGPDJEN1YFP_C_ (HIS3) or expressing pGPDYFP_N_ (URA3) or pGPDYFP_C_ (URA3), were used as controls.

## Acknowledgments

We thank Olivier Vincent and Bruno André for fruitful discussions. This work was supported by the strategic programme UID/BIA/04050/2013 (POCI-01-0145-FEDER-007569) and the project PTDC/BIA-MIC/5246/2020 funded by national funds through the FCT I.P. and by the ERDF through the COMPETE 2020 - Programa Operacional Competitividade e Internacionalização (POCI). GD was supported by “Fondation Sante”. CBA and GT acknowledge FCT for the PD/BD/135208/2017 and SFRH/BD/86221/2012 PhD grants, respectively.

